# Spt6 is a maintenance factor for centromeric CENP-A

**DOI:** 10.1101/560300

**Authors:** Georg OM Bobkov, Anming Huang, Sebastiaan J.W. van den Berg, Sreyoshi Mitra, Eduard Anselm, Vasiliki Lazou, Sarah Schunter, Regina Federle, Axel Imhof, Alexandra Lusser, Lars E.T. Jansen, Patrick Heun

## Abstract

Replication and transcription of genomic DNA requires partial disassembly of nucleosomes to allow progression of polymerases. This constitutes both an opportunity to remodel the underlying chromatin as well as the potential danger of losing epigenetic information. Centromeric transcription has been shown to be required for stable incorporation of the centromere-specific histone dCENP-A in M/G1-phase, which depends on the eviction of previously deposited H3/H3.3-placeholder nucleosomes. Here we demonstrate that the histone chaperone and transcription elongation factor Spt6 spatially and temporarily coincides with centromeric transcription and prevents the loss of old CENP-A nucleosomes in both *Drosophila* and human cells. Spt6 binds directly to dCENP-A and shows enhanced association with non-phosphorylatable dCENP-A mutants compared to histone H3, while phosphomimetic residues alleviate association with Spt6. We conclude that Spt6 acts as a conserved CENP-A maintenance factor, which is required during transcription-mediated chromatin remodelling at the centromere to ensure long-term stability of epigenetic centromere identity.

## Introduction

Centromeres constitute a platform for the assembly of the kinetochore during mitosis and mediate the attachment of chromosomes to the mitotic spindle for proper segregation of chromosomes. The position of the centromere is mostly determined epigenetically through the incorporation of the H3-variant CENP-A (also CID or dCENP-A in *Drosophila*) (Earnshaw & Rothfield 1985; Henikoff et al. 2000). Centromeric chromatin is composed of interspersed arrays of CENP-A and canonical histone H3 nucleosomes (Blower et al. 2002; Martins et al. 2016; Bergmann et al. 2011). While canonical H3 is replenished during DNA replication in S-phase (Ahmad & Henikoff 2002; Ahmad & Henikoff 2003), loading of CENP-A in *Drosophila*, humans as well as other vertebrates and takes place in a replication-independent manner from late mitosis to G1 (Jansen et al. 2007; Hemmerich et al. 2008; Bernad et al. 2011; Moree et al. 2011; Dunleavy et al. 2012; Silva et al. 2012; Lidsky et al. 2013). This process requires the exchange or removal of so-called ‘placeholder’ nucleosomes containing H3 and H3.3, which have been positioned on centromeric DNA-sequences during the previous S-phase (Dunleavy et al. 2011).

As expected for an epigenetic mark, centromeric CENP-A nucleosomes are remarkably stable and can be propagated not only over multiple cell divisions but also across generations. Indeed, epitope-tag labelling of dCENP-A revealed that once fully incorporated, the only turnover CENP-A nucleosomes experience in healthy proliferating cells is through replicative dilution (Bodor et al. 2013). Some of this stability is conferred to CENP-A by other centromere factors that act on the intact DNA-bound nucleosome itself. While CENP-C reshapes and clamps down the CENP-A nucleosome, CENP-N helps fastening CENP-A to the underlying DNA (Falk et al. 2015; Guo et al. 2017). Moreover, CENP-A nucleosomes that are assembled in mouse oocytes before birth, persist in the chromatin of prophase I-arrested cells for over a year and are sufficient for genome transmission to embryos through the entire fertile lifespan of the mouse (Smoak et al. 2016).

In actively dividing cells, however, chromatin is a highly dynamic structure. Cellular processes that require direct DNA contact like DNA replication or transcription induce large-scale chromatin remodelling events to allow the progression of DNA- and RNA-polymerases. This involves partial or full disassembly of nucleosomes (Ransom et al. 2010; Kulaeva et al. 2013), which challenges the stable transmission of epigenetic marks encoded in histone variants or histone tail modifications. Accordingly, mechanisms need to be in place that ensure the faithful transmission of epigenetic signals during replication and transcription.

Indeed, CENP-A nucleosomes are stably maintained during the replication of centromeric DNA (Bodor et al. 2013; Jansen et al. 2007; Mellone et al. 2011). Recent work identified the MCM2-7 replicative helicase to recycle previously deposited H3/H4, H3.3/H4 and CENP-A/H4 tetramers together with other chaperones (Alabert & Groth 2012; Burgess & Zhang 2013; Foltman et al. 2013; Gérard et al. 2006; Huang et al. 2015; Petryk et al. 2018). This ensures the transfer of parental nucleosomes to freshly replicated DNA and in human cells also includes the CENP-A specific chaperon HJURP previously only assigned to CENP-A loading (Zasadzińska et al. 2018).

Centromeres are also sites of active transcription, as revealed by the centromeric presence of RNA Polymerase II (RNAPII), centromeric RNA transcripts and transcription-associated histone modifications in various organisms including yeast, flies and humans (Catania et al. 2015; Choi et al. 2011; Bergmann et al. 2011; Bobkov et al. 2018; Chan et al. 2012; Eymery et al. 2009; Horard et al. 2009; Rošić et al. 2014; Quénet & Dalal 2014; Ohzeki et al. 2012; Sullivan & Karpen 2004). Centromeric transcription is important for centromere function (Ohkuni & Kitagawa 2011; Bergmann et al. 2011; Cardinale et al. 2009; Chen et al. 2015; Nakano et al. 2008), and it has been proposed that transcription-mediated chromatin remodelling plays a role in CENP-A loading (Choi et al. 2011; Chen et al. 2015; Bobkov et al. 2018). However, it is currently unclear how old CENP-A nucleosomes survive the passage of the elongating RNAPII. Active removal of CENP-A following transcription of centromeres has been reported in several organisms, albeit mostly following strong transcriptional upregulation induced experimentally (Bergmann et al. 2012; Hill & Bloom 1987). Moreover, persistent genotoxic stress correlates with activated centromeric transcription and a corresponding loss of CENP-A, similar to what is observed in permanently arrested senescent cells (Hédouin et al. 2017).

Eviction of nucleosomes through transcription is not unique to CENP-A-containing ones, but instead concerns all nucleosomes. To ensure genome integrity and avoid cryptic transcription, chromatin needs to be rapidly re-established in the wake of the DNA- and RNA polymerase. Whereas this is achieved during DNA replication through deposition of canonical histones, nucleosome gaps created by genomic transcription are filled through the replication-independent incorporation of H3.3 (Ahmad & Henikoff 2002a; Ahmad & Henikoff 2003; Ray-Gallet et al. 2011) as well as the recycling of displaced old histones. Disassembly of nucleosomes in front of a progressing RNAPII involves the histone chaperone <u>Fa</u>cilitates <u>C</u>hromatin <u>T</u>ranscription (FACT) (Foltman 2013; Kulaeva et al. 2010). FACT also acts to reassemble nucleosomes behind RNAPII together with the transcription elongation factor and histone chaperone Spt6. Spt6 can interact with histones, assembles them into nucleosomes (Bortvin & Winston 1996), and is able to increase the elongation rate of RNAPII both *in vitro* and *in vivo* (Endoh et al. 2004; Ardehali et al. 2009).

While a role for FACT at the centromere and its importance for CENP-A deposition has already been demonstrated in numerous organism (Foltz et al. 2006; Izuta et al. 2006; Okada et al. 2009; Chen et al. 2015; Choi et al. 2012; Prendergast et al. 2016), little is known about a centromeric function of Spt6. In budding yeast and flies, Spt6 was detected in a CENP-A pull-down and mass-spectrometry experiment (Barth et al. 2014; Ranjitkar et al. 2010). Mutants of Spt6 in budding yeast show segregation defects for a chromosome fragment (Basrai et al. 1996), whereas mutants in *S. pombe* exhibit CENP-A misincorporation genome-wide (Choi et al. 2012). Importantly, Spt6 prevents transcription-coupled loss of nucleosomes in gene bodies by reincorporating H3/H4 tetramers displaced during transcription. Consequently, Spt6 preserves the epigenetic information encoded in histone tail posttranslational modifications (PTMs) of recycled nucleosomes (Kato et al. 2013). Spt6 further stabilizes nucleosomes through a self-enforcing protein network comprising Spt6, the histone deacetylase Rpd3 and the H3K36 methyl-transferase Set2 by removing transcription-induced histone acetylation (Yoh et al. 2008; Drouin et al. 2010; Govind et al. 2010; Burugula et al. 2014). Consistent with a major role of Spt6 in the restoration of transcriptionally-perturbed chromatin in the wake of a progressing RNAPII, cryptic promoters are activated within transcription units in mutants for Spt6 (Kaplan et al. 2003; DeGennaro et al. 2014).

Here we demonstrate that transcription at the centromere, while being important for loading of new CENP-A, indeed poses a threat to the maintenance of ancestral CENP-A nucleosomes. However, long-term stability of the centromeric mark is achieved through effective recycling of expelled dCENP-A by Spt6 in both *Drosophila* and in human cells.

## Results

### Spt6 is present at mitotic and G1 centromeres

To identify novel factors associated with *Drosophila* centromeres we previously affinity-purified GFP-tagged dCENP-A containing nucleosomes and combined it with mass-spectroscopy analysis. Among the proteins enriched in dCENP-A containing chromatin was the transcription elongation factor and histone chaperone Spt6 (Barth et al. 2015).

To verify the identification of Spt6 as a centromere-associated protein, we investigated its cellular localization using fluorescent microscopy. Both endogenous Spt6 as well as the GFP-tagged transgene were detected at mitotic centromeres as well as in directly adjacent areas (Fig. 1a, b; upper panel). In interphase, Spt6 prominently stained euchromatic areas whereas the centromere surrounding heterochromatin was largely devoid of any signal. In a subpopulation of interphase cells, additional Spt6 foci that overlapped with centromere counterstaining were visible (Fig. 1a, b; lower panel). To characterize the localization of Spt6 with respect to various cell cycle stages in greater detail, we investigated cells that simultaneously expressed dCENP-A-mCherry and Spt6-GFP by live cell microscopy. In late G2 cells just prior to the entry into mitosis, Spt6-GFP was not detectable at centromeres (Fig. 1c and Supplementary Fig. 1a). While high levels of nucleo-cytoplasmic Spt6-GFP during mitosis interfered with its precise localisation *in vivo*, Spt6-GFP appeared at centromeres within minutes of entering the subsequent G1 phase (Fig.1c) and remained there for a period of 3-6 hours (Fig. S1a). Analysis of fixed S-phase cells labelled with 5-ethynyl-2′-deoxyuridine EdU (click-IT®) detected very little to no centromeric Spt6 (Fig. 1d, e), whereas investigation of midbody containing cells confirmed its presence at early G1 centromeres (Supplementary Fig. 1b).

**Fig. 1.**
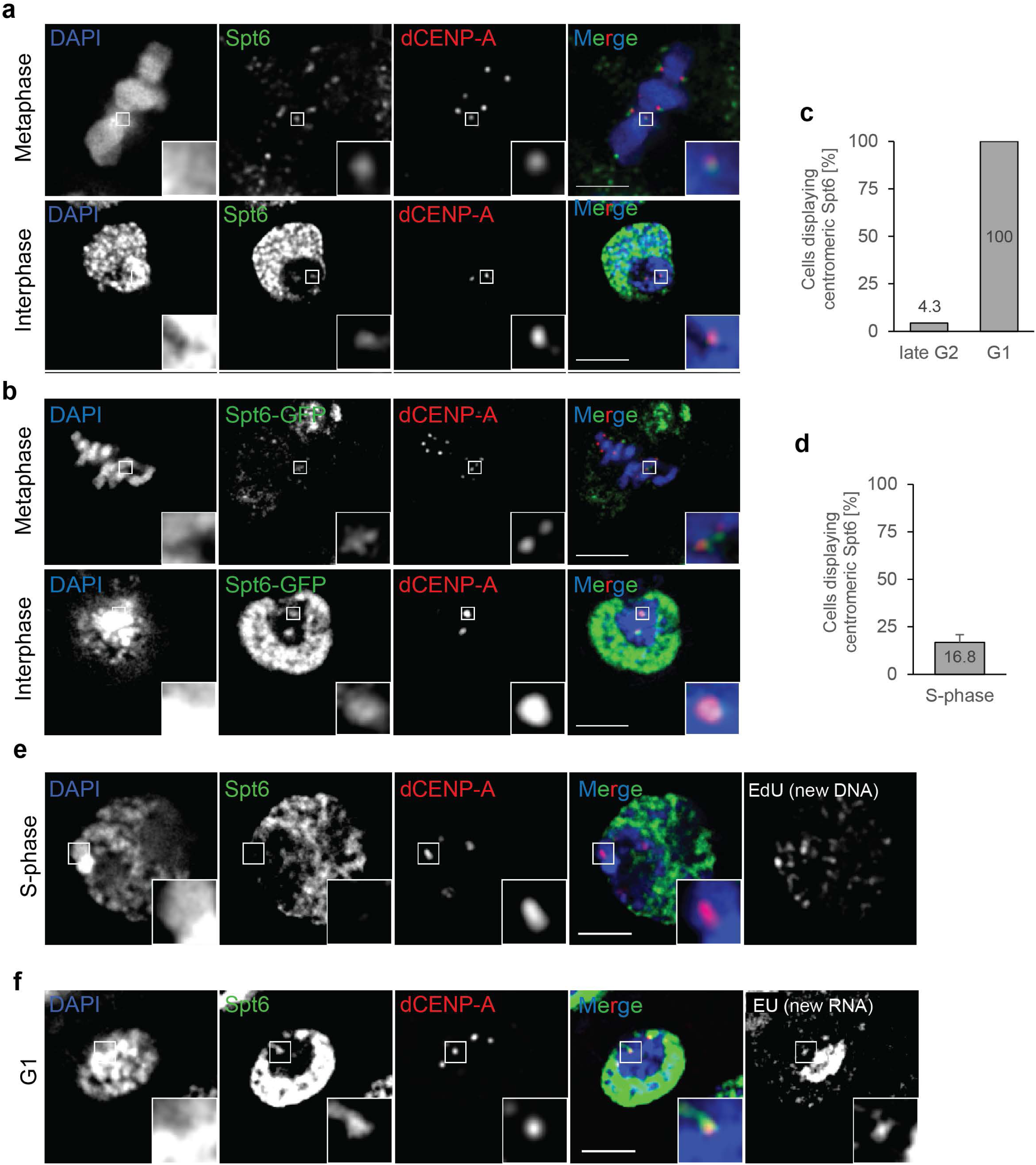
Spt6 localizes to centromeres in mitosis and G1. **a** Single optical section of fixed metaphase (upper panel) and interphase (lower panel) cells immunostained for endogenous Spt6 and dCENP-A. Boxes indicate the 3x enlarged inset. Scale bar represents 3 µm. **b** Single optical section of fixed metaphase (upper panel) and interphase (lower panel) S2 cells expressing Spt6-GFP. dCENP-A immunodetection served as a marker of centromeres. Boxes indicate the 3x enlarged inset. Scale bar represents 3 µm. **c** Analysis of the localization GFP-tagged Spt6 using life-imaging 1h before (late G2; n=23) and 1h after (G1; n=39) anaphase onset. Quantification was based on images shown in Supplementary Fig.1a. **d** Quantification of Spt6 localization in fixed EdU incorporating S-phase cells exemplified in Fig. 1e. N=3 replicates; n=30 cells. Data are represented as mean+SD. **e** Single optical section of fixed S-phase cell immunostained for Spt6 and dCENP-A. Incorporation of EdU served as a marker for S-phase. Boxes indicate the 3x enlarged inset. Scale bar represents 3 µm. **f** G1 phase S2 cell with nascent RNA production labelled by click-iT EU and immunostained for Spt6. Boxes indicate the 3x enlarged inset. Scale bars represent 3 µm.

This cell cycle-dependent localization pattern mirrors the previous mapping of centromeric RNAPII and centromere-associated transcripts to mitotic and G1 centromeres (Bobkov et al. 2018). Indeed, combining Spt6 immunostaining with pulse-labelling of nascent RNA using the Click-It® technology allowed the simultaneous detection of Spt6 together with centromere-associated transcripts at the same interphase centromeres (Fig. 1f). Taken together, centromeric association of Spt6 is cell cycle regulated and largely restricted to centromeres of mitotic and G1 cells.

### RNAi-mediated depletion of Spt6 leads to increased mitotic defects and an unspecific cell cycle arrest

Next, we decided to investigate the effects of Spt6 depletion via RNA interference (RNAi) in *Drosophila* S2 cells (Supplementary Fig. 2a). Interestingly, we observed a strong increase in mitotic defects in Spt6 depleted cells to levels comparable to prolonged depletion of dCENP-A (Supplementary Fig. 2b), mainly comprised of lagging chromosomes (Supplementary Fig. 2c). This suggests a potential impact on centromere functionality, which is consistent with previously reported missegregation of a chromosome fragment in a budding yeast Spt6 mutant (Basrai et al. 1996). However, during the course of the RNAi experiment, we noticed an apparent decrease of mitotic cells in samples treated for more than two days with dsRNAs targeting Spt6 (Supplementary Fig. 2d). Measurement of cell numbers in RNAi treated cultures confirmed that depletion of Spt6 leads to a cell cycle block whereas control cells depleted for the white protein were unaffected (Supplementary Fig. 2e). Subsequent FACS analysis revealed that the cell cycle arrest of Spt6 depleted cells was not specific and instead occurred across all cell cycle stages (Supplementary Fig. 2f).

### Centromeric dCENP-A levels are reduced following Spt6 degradation using an adapted deGradFP technique

Spt6 localization to centromeres in mitosis and G1 matches the time window where new dCENP-A is incorporated into *Drosophila* centromeres (Mellone et al. 2011; Dunleavy et al. 2012; Lidsky et al. 2013). Spt6 has further been shown to increase the elongation rate of RNAPII (M. Endoh et al. 2004; Ardehali et al. 2009) and recycle previously deposited nucleosomes during genomic transcription (Kato et al. 2013). CENP-A loading requires the exchange of placeholder nucleosomes (Dunleavy et al. 2011), which has been proposed to be mediated by transcription-induced chromatin remodelling (Choi et al. 2011; Chen et al. 2015; Bobkov et al. 2018). We therefore hypothesized that Spt6 might also play a role during centromeric transcription and the loading process of dCENP-A.

As loading of dCENP-A is coupled to progression through the cell cycle, the depletion of Spt6 via RNAi is not suitable to assess potential changes in dCENP-A deposition, especially as the cell cycle block already occurs with Spt6 levels only being reduced to roughly 50% (Supplementary Fig. 2a). To overcome this problem, we used the CRISPR/Cas9 technique to GFP-tag endogenous Spt6 between the second and third exon, inspired by the fully functional GFP-Spt6 protein produced in the GFP-TRAP study (Supplementary Fig. 3a) (Buszczak et al. 2007) and combined it with a degradation system targeting GFP (Caussinus et al. 2013). Western blot analysis confirmed that the GFP-fusion protein is the only form of Spt6 present in our stable S2 cell line (Clone C4, Supplementary Fig. 3b). Rapid, inducible degradation of GFP-Spt6 was achieved by adapting the deGradFP degradation system (Caussinus et al. 2013) to respond to the small molecule Shield1 (Banaszynski et al. 2006). Accordingly, we modified the original F-box-construct through addition of a degron domain (FKBP-L106P) that results in constant degradation of the fusion protein. Addition of Shield1 stabilizes FKBP-F-box-GFP Binding Protein (GBP), which then initiates the degradation of GFP-Spt6 (Fig. 2a). Indeed, we found that Shield1 induced rapid degradation of Spt6 within 12-24h as judged by IF of fixed cells and Western blot of total protein extracts (Fig. 2b, c). Similar to the RNAi-mediated depletion of Spt6, cell growth came to an arrest after 2 days of Shield1 treatment (data not shown). Interestingly, Shield1 treatment for 21h (< one generation time) led to a reduction of total dCENP-A at centromeres in Spt6-depleted cells (Fig. 2c, d), suggesting either impaired loading of new dCENP-A or increased loss of old dCENP-A.

**Fig. 2.**
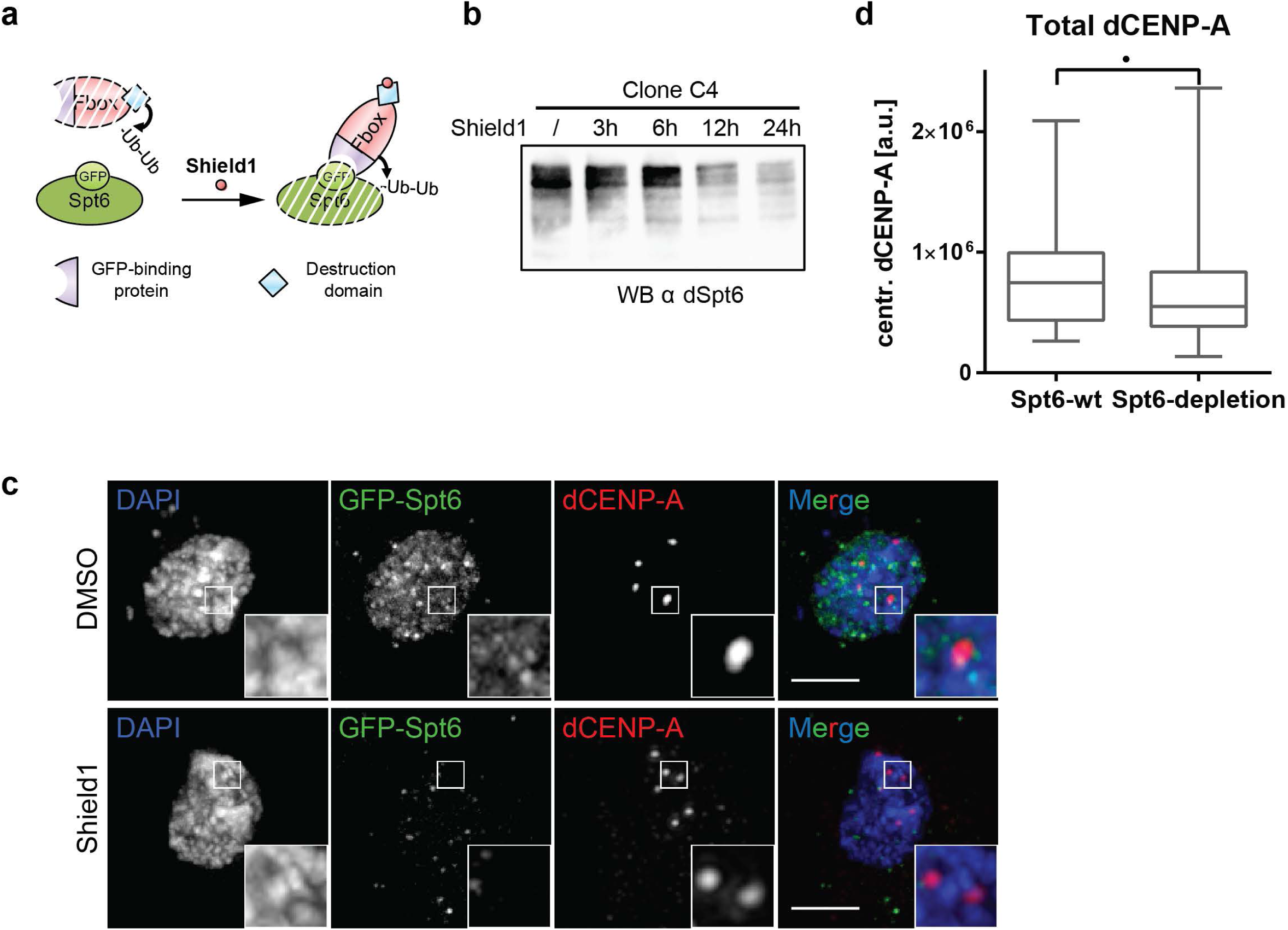
Spt6 depletion results in lower levels of total dCENP-A. **a** Schematic representation of the Shield1-induceable degradation of GFP-Spt6. **b** Western blot analysis demonstrating the depletion of GFP-tagged Spt6 following the addition of Shield1. **c** Maximum intensity projection of cells measured in Fig. 2d. Cells treated for 21h with DMSO (upper panel) or Shield1 (lower panel) are shown. Boxes indicate the 3x enlarged inset. Scale bar represents 3 µm. **d** Quantification of total centromeric dCENP-A levels in Spt6 wt or depleted cells. N=3 replicates; n=25 cells. Data are represented as mean+/-SD. • P≤0.05; •• P≤0.01; ••• P≤0.001 (unpaired t-test); n.s.=not significant.

### Spt6 prevents transcription-coupled loss of old dCENP-A

To distinguish between a defect in loading versus impaired maintenance of dCENP-A, we established the <u>R</u>ecombination Induced <u>T</u>ag <u>E</u>xchange (RITE)-technique (Verzijlbergen et al. 2010) in *Drosophila* S2 cells. This technique allows simultaneous tracking of both new (MYC-tagged) and old proteins (V5-tagged) via a Cre recombinase mediated epitope-tag switch (Fig. 3a). dCENP-A dynamics were assessed after a 21h treatment with Cre recombinase in the presence or absence of GFP-Spt6 (Fig. 3b). Levels of newly loaded dCENP-A^MYC^ were not significantly altered between both treatment conditions, although they showed a tendency towards increased loading in Spt6 depleted cells (Fig. 3c and 3e). In contrast, old dCENP-A^V5^ was significantly depleted from centromeres in cells where Spt6 was degraded (Fig. 3d, e).

**Fig. 3.**
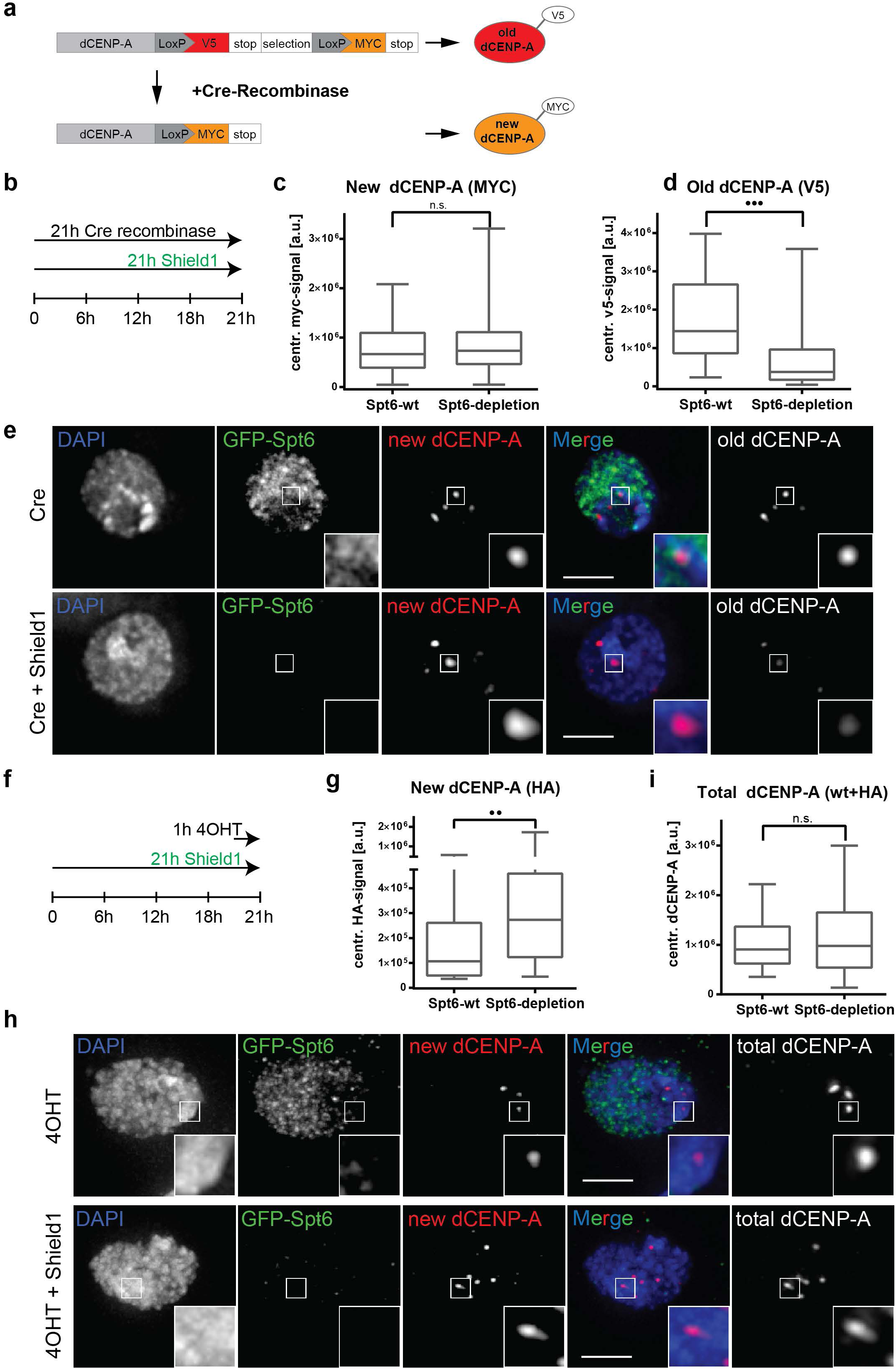
Parental dCENP-A is lost in Spt6 depleted cells. **a** Scheme displaying the RITE system used to distinguish old and new protein simultaneously (adapted from Verzijlbergen et al. 2010). **b** Experimental setup used in Fig. 3 c-e. **c**-**d** Quantification of the centromeric incorporation of new dCENP-A^MYC^ (**c)** or remaining levels of old dCENP-A^V5^ (**d)** at centromeres under Spt6 depletion conditions. N=3 replicates; n=10 cells. Data are represented as mean+/− SD. **e** Maximum intensity projection of cells measured in c-d. Cells treated with Cre recombinase (upper panel) or Cre recombinase plus Shield1 (lower panel) are shown. Boxes indicate the 4x enlarged inset. Scale bar represents 3 µm. **f** Experimental setup used in Fig. 3g-i. **g** Quantification of new centromeric dCENP-A^HA^ in fixed cells following 4OHT mediated release of dCENP-A_HA_ERT2 and depletion of Spt6 through Shield1 treatment. N=3 replicates, n=15 cells. Data are represented as mean+/− SD. **h** Maximum intensity projection of cells measured in Fig. 3g and 3i. 4OHT treated (upper panel) and 4OHT plus Shield1 treated cells (lower panel) are shown. Boxes indicate the 4x enlarged inset. Scale bar represents 3µm. **i** Quantification of total centromeric dCENP-A in fixed cells following 4OHT mediated release of dCENP-A_HA_ERT2 and depletion of Spt6 through Shield1 treatment. N=3 replicates, n=15 cells. Data are represented as mean+/− SD. •• P≤0.01; ••• P≤0.001 (unpaired t-test); n.s.=not significant.

Interestingly, the loss of old dCENP-A^V5^ was more pronounced than the reduction observed for total dCENP-A levels in Spt6 depleted cells (compare Fig. 2d and Fig. 3d), suggesting a partial compensation by increased incorporation of new dCENP-A. However, the RITE system did not reveal a significant increase in new CENP-A^MYC^, possibly because not enough new protein was produced during the course of the experiment to compensate for the loss of old dCENP-A^V5^. To test this hypothesis, we combined Spt6 degradation with a previously established tamoxifen-inducible HA-tagged dCENP-A (TI-dCENP-A^HA^) loading system (Bobkov et al. 2018). In this system, TI-dCENP-A^HA^ is constitutively produced but cannot participate in dCENP-A loading, as it is sequestered away in the cytoplasm due to an interaction with Hsp90. Only upon treatment with 4-hydroxytamoxifen (4OHT) is TI-dCENP-A^HA^ released and allows a larger pool of preproduced new dCENP-A proteins to become instantaneously available for incorporation into centromeric chromatin. Interestingly, when combined with simultaneous depletion of Spt6 using our inducible deGradFP system (Fig. 3f), we indeed observed a clear increase in incorporation of new dCENP-A compared to cells with wildtype levels of Spt6 (Fig. 3g, h). Moreover, the presence of endogenous dCENP-A combined with additional TI-dCENP-A^HA^ completely equalized total dCENP-A levels between cells with and without Spt6 depletion, in contrast to the reduction previously observed following Spt6 degradation (compare Figs. 2d and 3i).

### Depletion of human SPT6 leads to loss of human CENP-A maintenance

Next, we determined whether our observations are generalizable across species by depletion of the highly conserved SPT6 protein from human HeLa cells. We measured the rate of CENP-A retention across the cell cycle using SNAP-based fluorescent pulse labelling that specifically tracks old chromatin-bound CENP-A (Bodor et al. 2013; Bodor et al. 2012). As in *Drosophila* S2 cells, strong depletion of Spt6 results in a cell cycle arrest and secondary phenotypes. To prevent this, we aimed to only partially deplete Spt6 using specific siRNAs and controlled cell cycle progression by a thymidine arrest and release protocol (Fig. 4a, d). This hypomorphic setup enabled Spt6 depleted cells to pass through mitosis into G1, which was verified by monitoring Cyclin B levels (data not shown) and thus allow for a CENP-A loading event (Fig. 4a). Under these conditions, we observe a reproducible 20% reduction in CENP-A maintenance within a single cell cycle (Fig. 4b, c). It is important to note that a modest effect is expected as Spt6 depletion is, by necessity, incomplete. Nevertheless, we find these intermediate Spt6 levels to be rate limiting for CENP-A maintenance while they suffice for allowing cells to transit the cell cycle. CENP-C, previously identified as a major contributor to CENP-A maintenance (Falk et al. 2015) was used as a positive control (Fig. 4b, c), to demonstrate the maximum possible loss of CENP-A in these assays (~50%). In summary, these findings support a role for SPT6 in CENP-A maintenance in human cells, similar to our observations in *Drosophila*.

**Fig. 4.**
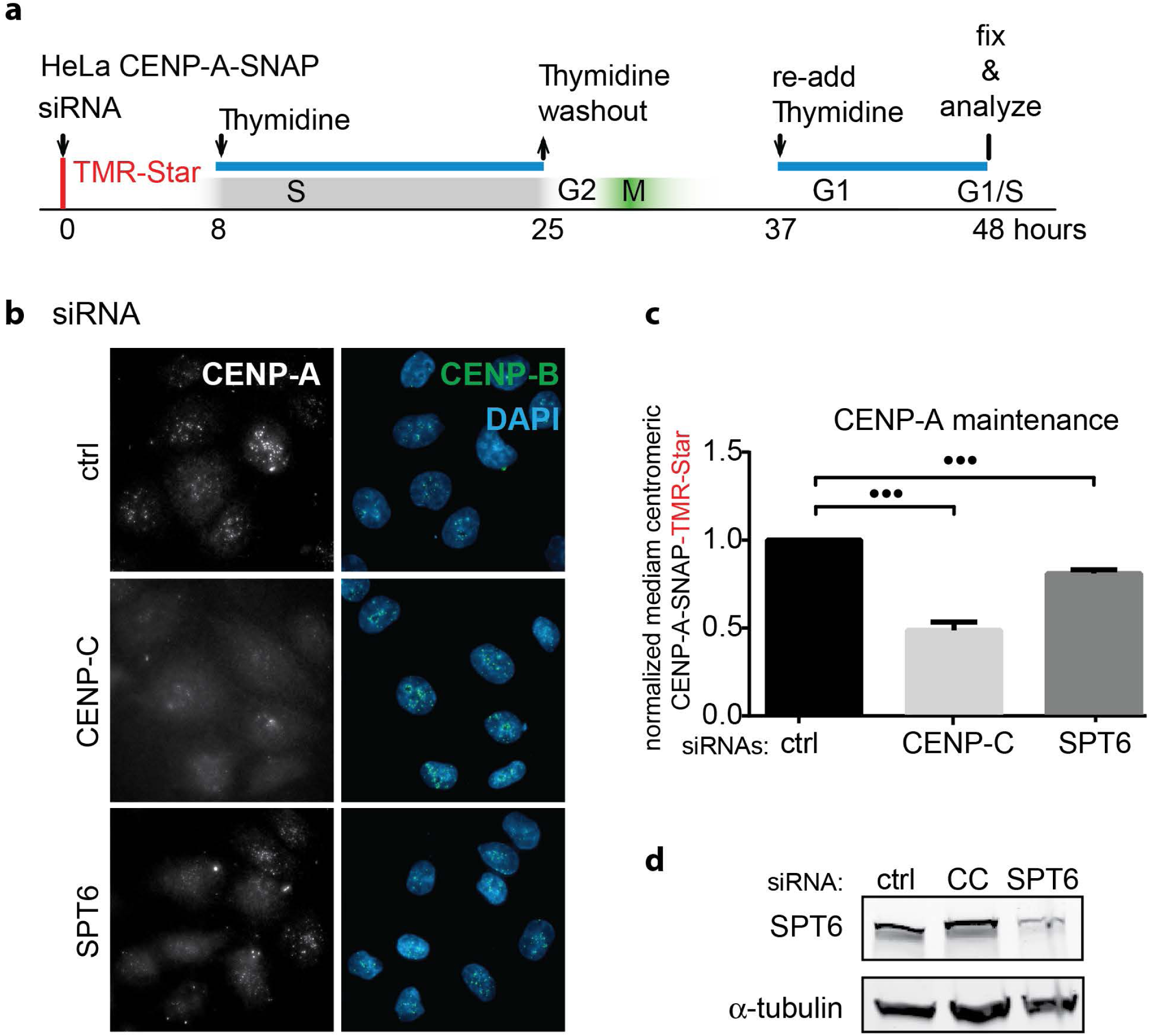
Depletion of human SPT6 leads to loss of CENP-A maintenance. **a** Experimental setup. HeLa cells expressing SNAP-tagged CENP-A were treated with TMR-star to detect previously incorporated CENP-A and siRNA treated to deplete indicated proteins. Cells were then synchronized in S phase by a thymidine block and release. Cells were allowed transit through G1 phase and were collected at the next G1/S boundary by re-addition of thymine. **b** Representative images of siRNA treated cells, 48 hours after TMR pulse labelling and mRNA depletion. Cells were counter stained with DAPI and anti-CENP-B antibodies to label DNA and centromeres, respectively. **c** Quantification of experiments shown in a and b. Mean +/− SD of 3 independent experiments is shown normalized to median control siRNA (ctrl). ••• P≤0.001 (unpaired t-test). **d** Cells were treated with indicated siRNAs for 48 hours and lysates were processed for immunoblotting and probed with indicated antibodies (CC = human CENP-C).

### dCENP-A binding to Spt6 and centromere abundance are affected by mutating phospho-residues

Intriguingly, the specific loss of previously deposited dCENP-A during the loading process of new dCENP-A in Spt6-depleted cells suggests that Spt6 binds and reincorporates not only H3 (Kato et al. 2013), but also dCENP-A/H4-tetramers. An association of Spt6 with dCENP-A might be expected due to our previously published mass spectroscopy of dCENP-A interactors (Barth et al. 2014). Although the exact binding interface is not known, binding of basic histones has been demonstrated for the unstructured highly acidic N-terminus of Spt6 in *S. cerevisiae* (Fig. 5a) (McDonald et al. 2010). To test whether the interaction between Spt6 and dCENP-A is direct, we expressed the N-terminal region of *Drosophila* Spt6 (residues 199-338) that encompasses the histone binding domain of yeast Spt6 (residues 239-314) based on amino acid sequence alignments (Supplementary Fig. 3c).

**Fig. 5.**
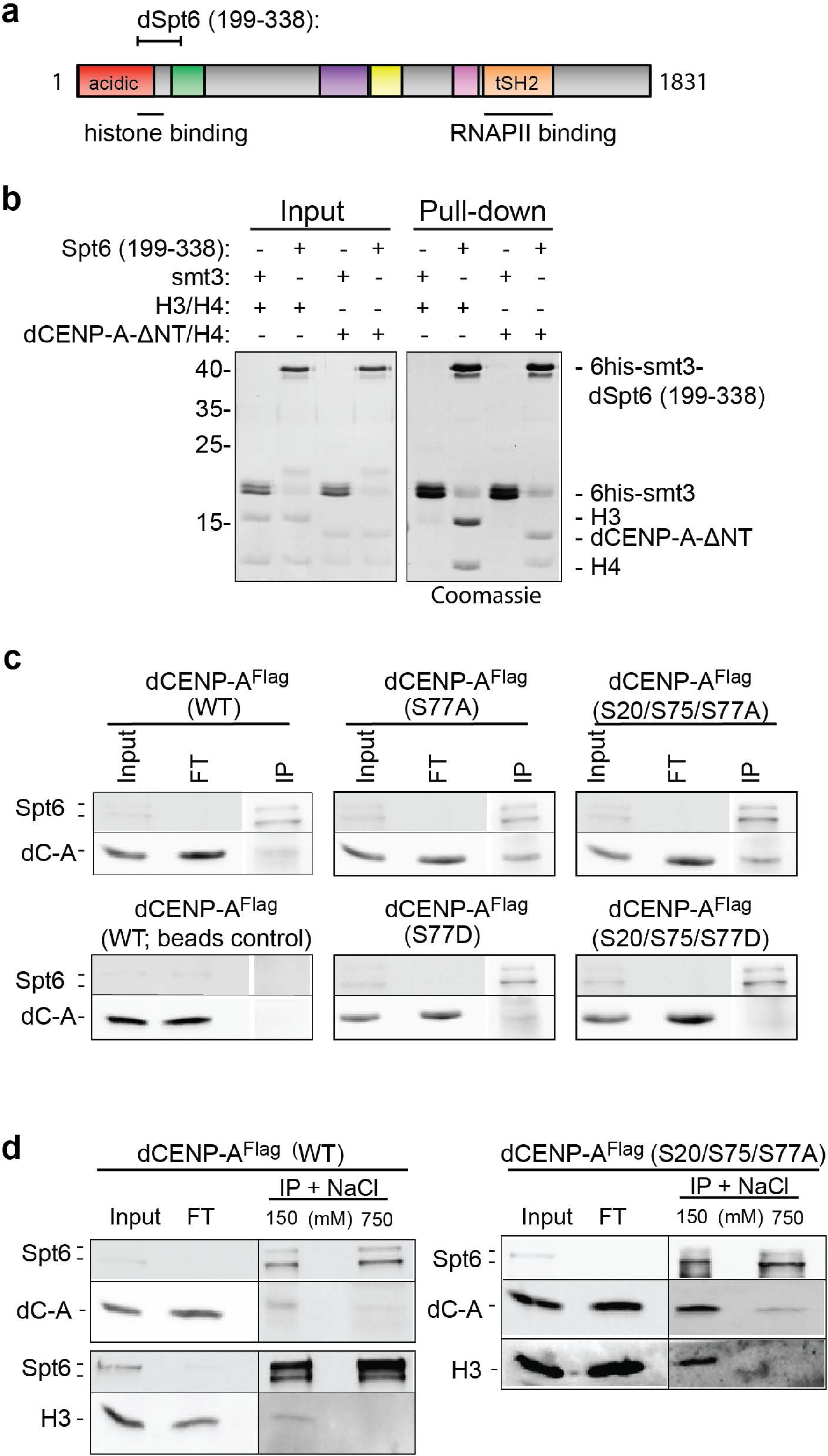
dCENP-A binds directly to Spt6 and is affected by mutating phosphoresidues**. a** *Drosophila* Spt6 domain organisation: acidic (red), Helix-turn-Helix (green), YqgF/RNaseH-like domain (purple), Helix-hairpin-Helix (yellow), S1 RNA-binding domain (magenta), tandem SH2 (orange). Corresponding histone and RNAPII binding domains based on *S. cerevisiae* (McDonald et al. 2010; Sdano et al. 2017). **b** Pull-down experiments of purified recombinant 6his-smt3-Spt6 (199-338) or 6his-smt3 as a negative control with recombinant H3/H4 or dCENP-A-ΔNT (101-225) are shown on a Coomassie-stained SDS-PAGE. **c** Western blot showing co-IPs of Spt6 with WT dCENP-A^FLAG^ and dCENP-A^FLAG^ bearing phosphorylation-abolishing (S to A) or phosphomimetic (S to D) mutations at phosphorylation sites S20, S75 and S77. FT, flowthrough; IP, immunoprecipitate. dC-A, dCENP-A. **d** Western blot showing co-IPs of endogenous Spt6 (two bands) with dCENP-A^FLAG^ (left) or dCENP-A^FLAG^ (S20/75/77A) (right) and H3 exposed to low (150 mM) and high (750 mM) salt wash conditions.

Recombinant Spt6 (199-338) was expressed and purified from bacteria and pull-down experiments revealed direct binding of Spt6 to both recombinant H3/H4 and dCENP-A-ΔNterm (101-255)/H4 tetramers, but not to the epitope tag alone control (6his-smt3) (Fig. 5b). As we could only produce soluble dCENP-A without its N-terminal tail in bacteria, we further confirmed that endogenous Spt6 is able to co-IP with full length dCENP-A^FLAG^ using *Drosophila* S2 cell extracts (Fig. 5c).

It has been shown that upregulated centromeric transcription in stressed and senescent murine cells can induce disassembly of CENP-A chromatin (Hédouin et al. 2017). However, simultaneous treatment with kinase inhibitors prevented the removal of CENP-A without affecting transcriptional upregulation, suggesting that phosphorylation events are required to achieve transcription-induced loss of CENP-A. As Spt6 prevents transcription-coupled loss of nucleosomes in gene bodies (Kato et al. 2013), this prompted us to generate cell lines that express Flag-tagged dCENP-A bearing mutations at three previously reported phosphorylation sites of its N-terminal tail (S20, S75 and S77; Boltengagen et al., 2016). The respective serines were mutated to either phosphomimetic aspartates or to non-phosphorylatable alanines. In line with a potential negative effect on dCENP-A retention through phosphorylation, all serine-to-alanine mutations increased the amount of dCENP-A interacting with Spt6 when compared to wild-type dCENP-A. In contrast, a mutation to aspartate at S77 weakened and triple aspartate substitution at S20, S75 and S77 completely abolished co-immunoprecipitation of dCENP-A with Spt6 (Fig. 5c). To test whether Spt6 shows different interaction specificity for H3 and dCENP-A, we exposed the co-IP’s to high salt (750 mM) extraction. While high salt treatment strongly disrupted the association of Spt6 with canonical H3, a small subpopulation of dCENP-A^FLAG^ remained bound (Fig. 5d). Interestingly, the salt-resistant fraction became more pronounced when using the non-phosphorylatable triple alanine dCENP-A mutant (S20/S75/S77A) (Fig. 5d).

Next, we tested whether the altered interactions between Spt6 and dCENP-A-mutants might also correlate with changed dCENP-A abundance at the centromere. We therefore quantified centromeric signals of wild-type, S77A or S77D dCENP-A in S2 cells. Consistent with the co-IP results, dCENP-A-S77A abundance was significantly higher as compared to wild-type dCENP-A (Fig. 6a, b), while dCENP-A-S77D intensities were strongly decreased (Fig. 6c, d).

**Fig. 6.**
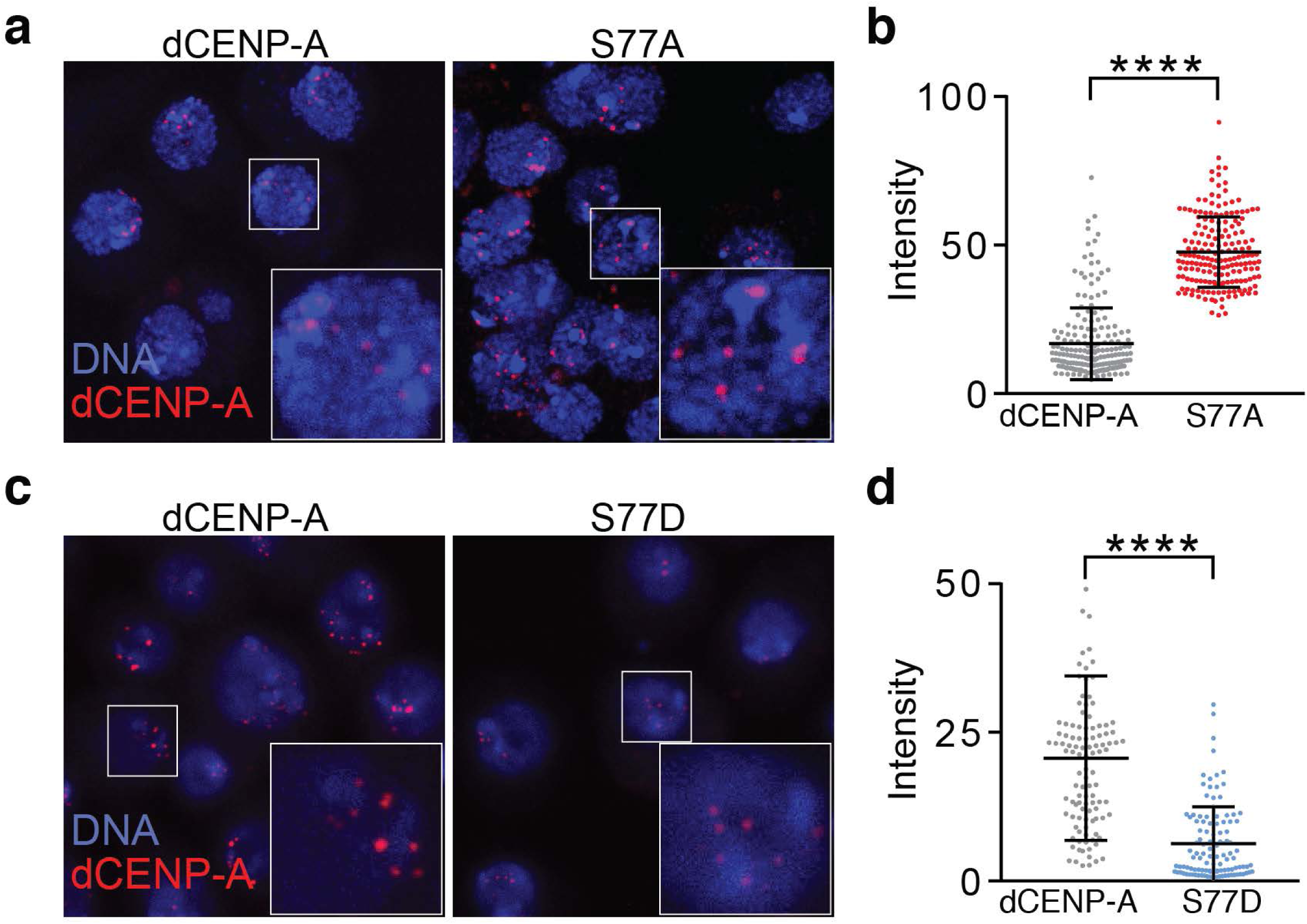
dCENP-A abundance at the centromere is affected by phosphorylation. **a** Stably transfected SNAP-tagged wild-type or S77A mutant dCENP-A was visualized by staining with TMR Star. Boxes indicate the 2.5 times enlarged inset. **b** Quantification of centromeric signal intensities of wild-type (n=182) and S77A CENP-A (n=274). Representative images (**a)** and quantification (**b)** of one out of three independent experiments are shown. **c** SNAP-tagged wild-type and S77D-mutant CENP-A staining by TMR Star. Boxes indicate the 2.5 times enlarged inset. **d** Quantification of centromeric signal intensities of wild-type (n=110) and S77D CENP-A (n=112). Representative images (**c)** and quantification (**d)** of one out of five independent experiments are shown. Statistical significance (threshold p<0.05) was determined by unpaired t-test (**** P<0.0001).

## Discussion

The CENP-A nucleosome is considered to be the key epigenetic mark for centromere identity in most organisms. Accordingly, CENP-A and epigenetic marks in general should meet three requirements: (1) Template its own deposition, (2) be replenished in a cell cycle-controlled manner to counteract dilution by half in each S-phase and (3) be stably transmitted to the next cell generation (Gómez-Rodríguez & Jansen 2013).

As such, new dCENP-A can be targeted to sites of previous CENP-A deposition by its chaperone CAL1, which is recruited to centromeres by dCENP-C (Chen et al. 2014; Erhardt et al. 2008; Olszak et al. 2011; Schittenhelm et al. 2010). Loading of new CENP-A is restricted to mitosis and G1 (Jansen et al. 2007; Mellone et al. 2011; Dunleavy et al. 2012; Lidsky et al. 2013) and serves primarily to replenish CENP-A containing nucleosomes that became diluted by half during the preceding S-phase. During DNA replication the MCM2-7 replicative helicase along with other histone chaperones like HJURP, are instrumental for the stable transmission of parental CENP-A during S-phase (Huang et al. 2015; Petryk et al. 2018; Zasadzińska et al. 2018).

We have recently shown in *Drosophila* S2 cells that transcription at the centromere is required for stable nucleosome incorporation of new dCENP-A (Bobkov et al. 2018). This finding could be explained by a model in which chromatin remodelling is re-purposed to evict placeholder H3 nucleosomes to make room for deposition of new dCENP-A. However, the induction of nucleosome eviction during CENP-A loading also bears the danger of losing previously incorporated CENP-A (Supplementary Fig. 4). Here we report the identification of the transcription elongation factor and histone chaperone Spt6 as a new CENP-A maintenance factor, which safeguards previously deposited CENP-A during centromeric transcription (Fig. 7).

**Fig. 7.**
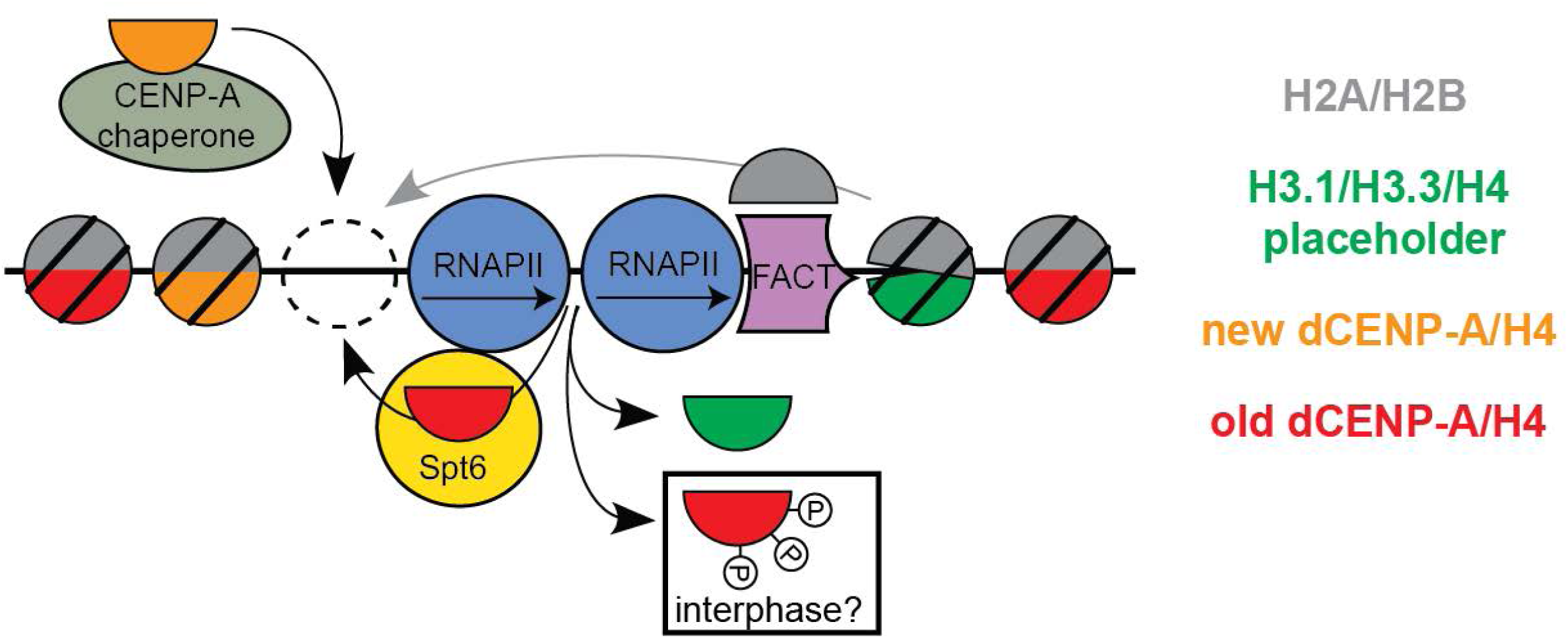
Model showing the histone dynamics during dCENP-A loading. Both old CENP-A and placeholder H3 nucleosomes can be evicted during the transcription-induced remodeling of centromeric chromatin, with Spt6 retaining old CENP-A nucleosomes. Phosphorylation of CENP-A interferes with Spt6-mediated recycling and maintenance.

We find that *Drosophila* Spt6 localizes to centromeres during mitosis and G1 (Fig. 1 and Supplementary Fig. 1), coinciding with the time window when transcription and dCENP-A loading occurs (Bobkov et al. 2018; Dunleavy et al. 2012; Lidsky et al. 2013; Mellone et al. 2011). The SH2 domain enables Spt6 to interact directly with RNAPII (Yoh et al. 2007; Close et al. 2011; Diebold et al. 2010; Liu et al. 2011; Sdano et al. 2017), therefore it is likely that recruitment of Spt6 to centromeres is a direct consequence of centromeric transcription during these cell cycle phases. Because Spt6 prevents transcription-coupled loss of posttranslationally modified nucleosomes in gene bodies (Kato et al. 2013), we tested whether Spt6 might act to maintain dCENP-A at the centromere. Indeed, when we depleted Spt6 in *Drosophila* or human cells, we observed a specific loss of old CENP-A after passage through mitosis into G1 phase (Fig. 3 and 4). This observation suggests that ongoing transcription evicts nucleosomes at centromeres and that Spt6 serves a conserved role to recycle CENP-A/H4 tetramers expelled by closely spaced polymerase complexes (Fig. 7) (Kulaeva et al. 2010). The model further predicts that the creation of nucleosomal gaps is a prerequisite for full incorporation of new dCENP-A. Consequently, the additional loss of nucleosomes in Spt6 depleted cells should create more opportunities to load new dCENP-A. In line with such a hypothesis, we observed a clear increase in loading of new TI-dCENP-A^HA^ (Fig. 3g, h), when we used an experimental system that provides elevated levels of ready-made TI-dCENP-A^H^ for loading (Fig. 3g, h). This is further supported by the fact that the expected loss of total centromeric dCENP-A in Spt6 depleted cells is completely compensated under these conditions (compare Fig. 2d and 3i).

We currently do not know if the mitotic defects observed upon Spt6 depletion by RNAi are a direct or indirect consequence of the removal of this protein (Supplementary Fig. 2b-d). It is possible that instant loss of old nucleosomes with specific PTMs at the centromere might have an immediate effect on chromosome segregation. PTMs important for centromere function have been identified on CENP-A and shown to affect CENP-A stability and correct mitotic progression (Bade et al. 2014; Bailey et al. 2013; Goutte-Gattat et al. 2013; Niikura et al. 2015; Samel et al. 2012). Moreover, methylation of lysine 20 on the associated H4 plays an essential role for kinetochore formation (Hori et al. 2014). Likewise, in addition to CENP-A nucleosomes, centromeres contain canonical H3 nucleosomes with a specific set of posttranslational modifications that might need to be retained (Sullivan & Karpen 2004; Bergmann et al. 2011). We therefore postulate that Spt6 should be able to distinguish between “placeholder” nucleosomes that need to be removed and epigenetically marked nucleosomes that should be kept. As previously demonstrated for H3/H4 in budding yeast (McDonald et al. 2010), we observe direct binding of a bacterially expressed N-terminal fragment of Spt6 (199-338) with both H3/H4 and dCENP-AΔNT/H4 tetramers (Fig. 5b). In addition, full length dCENP-A^FLAG^ and H3 co-IP with endogenous Spt6 from S2 cell extracts with comparable efficiency (Fig. 5d).

Interestingly, CENP-A is phosphorylated in various organism including flies and humans (Bailey et al. 2013; Boltengagen et al. 2016) and phosphorylation events have been linked to transcription-induced loss of centromeric CENP-A nucleosomes in mouse cells (Hédouin et al. 2017). To test whether phosphorylation of dCENP-A affects its maintenance, we mutated three previously identified phospho sites in the N-terminal tail of dCENP-A (S20, S75 and S77; Boltengagen et al., 2016). Indeed, we found that dCENP-A mutants carrying the phosphomimetic residue aspartate showed significantly reduced binding to Spt6, while the opposite was observed for the respective non-phosphorylatable alanine mutants (Fig.5c). Furthermore, non-phosphorylatable mutants of dCENP-A bound more robustly to Spt6 when exposed to high salt washes than canonical H3. This difference might enable Spt6 to distinguish between the two histone H3-variants and allow selective retention of CENP-A, while placeholder nucleosomes are exchanged. Consistent with the observations described above, centromeres of stably transfected S2 cells show lower abundance of a phosophomimetic and an increase of a non-phosphorylatable dCENP-A mutant as compared to wildtype dCENP-A.

Taken together, we propose that the transcription-mediated eviction of centromeric nucleosomes affects both placeholder H3 and previously deposited CENP-A nucleosomes. However, loss of the centromere mark is prevented by specific recycling of CENP-A potentially involving phospho-regulation of the CENP-A/Spt6 interaction (Fig. 3, 4 and 7). We conclude that Spt6 acts as an important CENP-A maintenance factor and contributes to the long-term stability of the epigenetic centromere mark.

## Methods

### Plasmid constructs

Spt6 was PCR’ed from genomic S2 cell cDNA and cloned into the SpeI/EcoRV sites of a pMT-GFP-hygro vector, creating pMT-Spt6-GFP-hygro. For the GFP-Spt6 IP, the SNAP tag was inserted C-terminally by PCR using Xho1/SacII into a pMT-CID-V5 vector (Heun et al. 2006). The CID-HA-ERT2 vector is described (Bobkov et al. 2018). The RITE-(V5/MYC) plasmid was created by combining synthesized g-block DNA fragments with a pMT-CID-V5 (Invitrogen) plasmid. The 1kup_resGFPSpt6_stop-plasmid was generated by combining the PCR’ed genomic region of Spt6 1kb upstream fragment and 2kb downstream fragment into a pIB-vector backbone (Invitrogen) using BstZ17I/NheI and KpnI/SacII respectively. A g-block synthesized artificial GFP-exon was cloned between the 2^nd^ and 3^rd^ Spt6-exon using AscI and SexA1.

To generate dCENP-A-S77A or -S77D single mutant constructs and the dCENP-A-S20/75/77A and CENP-A-S20/75/77D triple-mutant constructs site-directed mutagenesis was performed using the previously described SNAP-CENP-A pMT-puro or the Strep-Flag-CENP-A pMT-puro plasmid (Boltengagen et al., NAR 2016) as a template. Primer sequences are available upon request.

### Cell culture

*Drosophila* S2 Schneider cells were grown at 25°C in Schneider’s *Drosophila* medium (SERVA) supplemented with 10% fetal calf serum and antibiotics (0.3 mg/ml penicillin, 0.3 mg/ml streptomycin). Cells were transfected using the XtremeGENE HP transfection reagent (Roche) or Effectene Transfection Reagent (QIAGEN), and stable lines were selected with 100 μg/mL Hygromycin B, 2 μg/ml puromycin or 10 µg/ml puromycin for pMT-Puro transfection constructs. In the latter case, cells were maintained in 2 µg/ml puromycin after selection. The endogenously tagged GFP-Spt6 clone using CRISPR was transfected with pMT-CID-SNAP-V5-Hygro to create a stable cell line used in GFP-Spt6 IP.

HeLa CENP-A-SNAP cells were grown at 37°C, 5% CO_2_ in DMEM cell culture media supplemented with 10% newborn calf serum (NCS), 2 mM glutamine, 1 mM sodium pyruvate (SP), 100 U/ml penicillin, and 100 μg/ml streptomycin. The HeLa monoclonal cell lines stably expressing CENP-A-SNAP have been previously described (Jansen et al. 2007 clone #72).

### RNAi

Exponentially growing S2 cells were incubated for 30’ in 1 ml serum-free medium containing 20 µg of dsRNA before 3 ml of serum-containing medium were added. Cells were harvested after 3 (white/Spt6) or 7 (dCENP-A) days. Primers used for dsRNA synthesis:

Spt6_F: TTAATACGACTCACTATAGGGATGCCGGGCAAGTTCCTGCTGTCCTA

Spt6_R: TTAATACGACTCACTATAGGGGGCGTCATGAACGGAGTCTGTCCAC

dWhite_F: TTAATACGACTCACTATAGGGACTGCTCAATGGCCAACCTGTGGAC

dWhite_R: TTAATACGACTCACTATAGGGCCTCGGCCATCAGAAGGATCTTGTC

RNAi in HeLa cells was performed in a 24-well format with 2.5 pmol of small interfering RNAs final concentration (siRNAs) using Lipofectamine® RNAiMAX Transfection Reagent (Invitrogen) according to the manufacturer’s instructions. All siRNAs were obtained from Silencer® Select Pre-Designed & Validated siRNA (Life Technologies): CENP-C (s2913), SPT6 (s13634, s13635). Neg9 depletion siRNA target 5’-UACGACCGGUCUAUCGUAGTT −3′ was used as a control (ctrl). Following RNAi, HeLa cells were synchronized by a single thymidine block for 17h as described (Bodor et al. 2012), followed by a release into G2 and through G1 and are collected by re-addition of Thymidine, as outlined in Figure 4a.

### Immunofluorescence and SNAP labelling

Generally, S2 cells were settled for 20’ on polylysine coated cells and fixed for 7’ with 3.7% formaldehyde solution (Sigma) in PBS or in −20°C methanol. For staining of endogenous Spt6 at centromeres, shorter fixation conditions (3’ in PBS/0.1% Triton (PBS-T) containing 1.85% formaldehyde) was required. Following a wash in PBS-T, samples were blocked with Image-iT™FX signal enhancer (Invitrogen) for at least 30’. Unless otherwise noted, all antibodies were used 1:100 diluted: chicken α dCENP-A (1:20; P. Heun), rat α dCENP-A (4F8; E. Kremmer), rabbit α H3S10p (Abcam), mouse α dSpt6 (26D4 1:50; E. Kremmer), rat α dSpt6 (13D4 1:50; E. Kremmer), mouse α GFP (Clone 496; D. van Essen), mouse α tubulin (Sigma) and mouse α V5 (Invitrogen). Secondary antibodies coupled to Alexa 488, Alexa 555 and Alexa 647 fluorophores (Invitrogen) were used at a 1:100 dilution. Counterstaining of DNA was performed by DAPI (5 µg/ml; 3’). Antibodies against CENP-B (Santa Cruz Biotechnology, sc-22788 Rabbit polyclonal) and cyclin B1 (sc-245; Santa Cruz Biotechnology) were used at dilutions of 1:1000 and 1:50. Fluorescent secondary antibodies Donkey anti-Rabbit FITC and Donkey anti-mouse 680 (Rockland) were used at a dilution of 1:200. To visualize SNAP-tagged dCENP-A proteins cells were labelled with 3 µM SNAP-Cell TMR Star (New England Biolabs) for 30 min. Non-reacted TMR Star was washed out, cells were fixed in 3.7% paraformaldehyde/0.3% Triton-X100 for 12 min and washed in PBS. Nuclei were counterstained with DAPI (1 μg/ml; 10 min) and cells were mounted in vectashield (Vector Laboratories). SNAP labeling in HeLa cells was performed essentially as described (Bodor et al. 2012). Cells were labeled for 15 min with 2 μM TMR-Star (New England Biolabs) in complete medium for pulse labeling, after which cells were washed twice with phosphate-buffered saline (PBS) and reincubated with complete medium. After an additional 30 min, cells were washed once more with PBS and reincubated with complete medium with 2 μM BTP (SNAP-Cell Block; New England Biolabs) for 30 minutes and again washed with PBS and further treated for analysis, as indicated (Figure 4a). HeLa cell fixation, immunofluorescence, and 4′,6-diamidino-2-phenylindole (DAPI) staining was performed as described (Bodor et al. 2012), except for the secondary antibody incubation which was performed for 30 minutes at 37°C.

### Click-it chemistry

S-phase cells were visualized using the Click-iT® EdU Imaging Kit from Thermo Fisher Scientific following the manufacturer’s instructions (labelling: 15’/10 µM EdU). Global RNA transcription was detected using the Click-iT® RNA Imaging Kit from Thermo Fisher Scientific (labelling: 5’/4 mM EU). Cells were pelleted and resuspended twice in 1 ml of medium to allow unbound EU to diffuse (5’; 400 rpm), before cells were settled and fixed as usual. Click-it reaction was performed according to the manufacturer’s instruction.

### Microscopy and image analysis

All images (see exception below) were taken on a DeltaVision Elite Imaging System and were deconvolved using softWoRx Explorer Suite (Applied precision). Images of fixed cells were taken as 50-65z stacks of 0.2 µM increments using a 100x oil immersion objective. Time-lapse imaging was performed with 25z stacks of 0.4 µM increments using a 60x oil immersion objective and a time-lapse of 2’. Quantification of signal intensities were performed using the softWoRx Explorer Suite. Average background of five non-centromeric nuclear measurements was subtracted from measured centromeric signal (dCENP-A, HA, V5 and MYC analysis) and average background of five cytoplasmic measurements was subtracted from five measured nuclear signals (Spt6). Images of SNAP-tagged dCENP-A expressing S2 cells were taken on a Leica TCS SP5 instrument. Image acquisition was performed using a 63x oil objective with a pixel size of 48.1 nm and by collecting 0.13 µm z-sections spanning the entire nuclei. 3D images were reconstructed and analysed by Imaris V5.1. Mean TMR ± SEM intensities were calculated and statistical significance was determined separately for each experiment by t-test using GraphPad Prism 7.0 software. HeLa cells were imaged on a DeltaVision Core system (Applied Precision) and centromeres were quantified with CRaQ as described (Bodor et al. 2012).

### Whole cell lysates

All steps were carried out on ice/at 4°C and all used buffers contained in addition Protease inhibitor cocktail tablets (cOmplete Tablets; Roche) and 0.5 mM PMSF. Cells were washed twice with PBS before lysis. Pelleted cells were resuspended in buffer L (50 mM Tris-HCl, 150 mM NaCl, 1 mM EDTA, 1% Triton X-100, 1 mM MgCl_2_,) and sonicated 10x with an interval of 30’’ on “medium” setting (Bioruptor300; Diagenode). Protein concentrations were measured using Quick Start™ Bradford 1x Dye Reagent from BIO RAD according to the manufacturers manual and equal amounts loaded for each lane.

### Western Blot analysis

*Drosophila* S2 cell samples were boiled for 10’ in loading buffer separated on 10%-12% (fractionation assay), 6% (Spt6/RNAPII whole cell lysates) or 10% (Co-IPs) SDS-PAGE gels and processed for western blotting using mouse α GFP (1:2000, Clone 71; D. van Essen), mouse α Spt6 (25C3 1:500; E. Kremmer/A. Schepers), rabbit α CENP-A (1:5000, ab10887, Abcam), rabbit α H3 (1: 20000) and mouse α tubulin AA4.3 (1:1000; DSHB). Secondary antibodies coupled to horseradish peroxidase (Dianova) were used at 1:10000.

HeLa Cells were boiled in 2x Laemmli sample buffer and whole cell lysates were separated by SDS-PAGE and transferred onto nitrocellulose membranes for antibody incubation. Antibodies used were SPT6 (Abcam, ab32820 rabbit polyclonal) and α-tubulin (Sigma, T9026 mouse monoclonal). Secondary antibodies Donkey anti-Rabbit 800 (Li-Cor, 926-32211) and Donkey anti-mouse 680 (Rockland, 610-744-124) were used.

### Coimmunoprecipitation

All steps were carried out on ice/at 4°C, all used buffers contained in addition protease inhibitor cocktail tablets (cOmplete Tablets; Roche) and 0.5 mM PMSF. Mouse anti-Spt6 antibody (25C3; E. Kremmer/A. Schepers) was coupled to Protein A-Sepharose CL-4B (GE Healthcare) beads and stored at 4°C until actual pulldown. For nuclei purification, 50-100×10^6^ cells were pelleted (10’; 1000×g) and resuspended in 5 ml of nuclear buffer A (85 mM KCL, 5.5% sucrose, 10 mM Tris-HCl pH7.5, 0.2 mM EDTA, 0.5 mM spermidine). 5 ml of nuclear buffer B (as A but containing 0.5% NP40) was added and samples incubated for 3’ on ice. After centrifugation of nuclei (10’; 2000×g), supernatant was discarded, the pellet was washed twice with hypotonic buffer (20 mM HEPES-KOH pH7.9, 20 mM NaCl, 5 mM MgCl_2_, 10 mM imidazole and 0.5 mM β-mercaptoethanol) and resuspended in 300 µl of hypotonic buffer containing 0.5% NP-40 (Sigma-Aldrich) and 200 U Benzonase (Novagen) and incubated with rotation for 1h. NaCl was added to a final concentration of 300 mM and samples were rotated for another hour before salt concentration was brought to 150 mM through dilution with hypotonic buffer containing 0.5% NP-40 (Sigma-Aldrich). 10% of the supernatant after centrifugation (15’; 15000×g) served as input and the remaining sample was either split into two or three equal samples. Antibody-bound sepharose beads were added to each sample and rotated overnight. 10% of the unbound supernatant served as flowthrough and beads were washed for 1h with hypotonic buffer containing 0.5% NP-40 and either 150 mM or 750 mM NaCl (Fig. 5c). For Fig. 5d, washes were performed with 150 mM salt buffer. 10% of each supernatant served as the wash sample, beads were washed three additional times for 2’ with buffer containing the respective salt concentration before bound proteins were eluted by addition of 2× Laemmli sample buffer to obtain the IP sample.

### Expression of recombinant proteins and pull-down assay

6his-smt3-Spt6 (199-338) and 6-his-smt3 were recombinantly expressed in bacteria. Histones H3/H4 and CENP-A-ΔNT (101-225)/H4 were purified as described in (Voigt et al. 2012), 6his-smt3-Spt6 (199-338) and 6his-smt3 were incubated with dCENP-A-ΔNT(101-225) or H3/H4 in Pull-Down Buffer containing 20 mM Tris-HCl pH 8, 0.5 mM β-Mercaptoethanol, 10 mM imidazole, 0.1% Triton X-100, 5% glycerol and a final concentration of 250 mM NaCl on an overhead rotator for 1h at 4°C. Afterwards, input was taken and 20 µl HIS-Select® HF Nickel Affinity Gel (Sigma) pre-equilibrated with Pull-Down Buffer was added to each sample and incubated on an overhead rotator for 1h at 4°C. Beads were washed 5x with 1 ml Pull-Down Buffer, bound proteins eluted with Laemmli Buffer at 95°C for 5 min and analysed on 15% SDS-PAGE gel stained with InstantBlue™ (Expedeon).

### CRISPR/Cas9

S2 cells were transfected with a plasmid containing pIB_Cas9_CD4_Blast, the plasmid providing the GFP-tagged template (1kup_resGFPSpt6_stop) and two PCR products delivering the guide RNA’s. Guide RNA sequences were selected using the flycrispr-tool (http://tools.flycrispr.molbio.wisc.edu/targetfinder/; sgRNA1: TGACGACTTCTCAAAGTACGAGG; sgRNA2: GGTTACGATTCCGATGGCGTCGG). After 2 d, transfected cells were selected for 4 d with blasticidin. GFP-positive cells were picked under the microscope and clonally amplified.

### Cell cycle analysis

Cells were pelleted in a FACS tube (7’, 1000xg) and fixed at 4°C overnight in 70% ethanol. Fixed cells were stained for 1h in the dark (50 µg/ml propidium iodide, 100 µg/ml RNase in PBS) and directly subjected to analysis on a BD FACScalibur Flow Cytometer using a gate for single cells.

---

**Supplemental Fig. 1.**
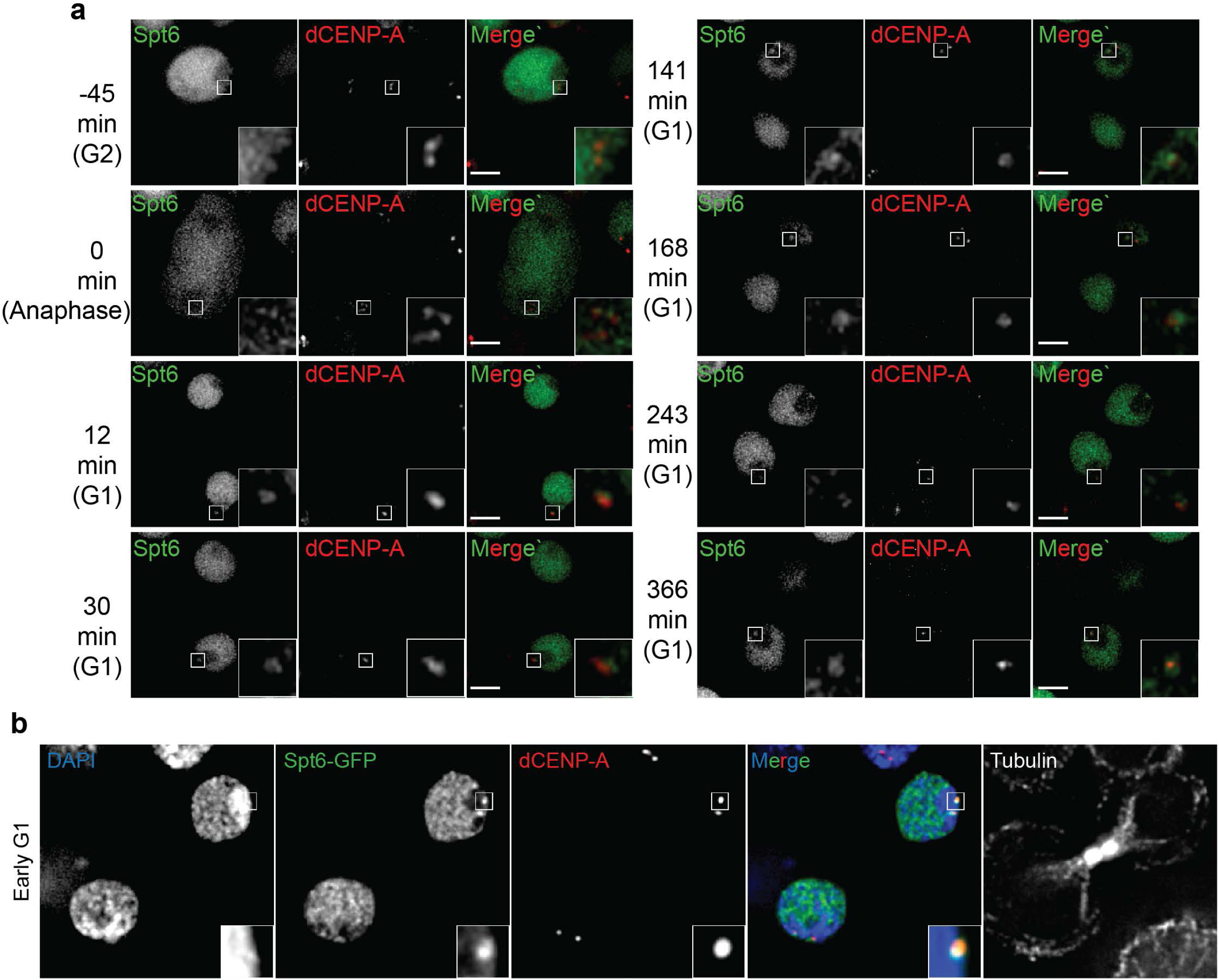
Spt6 localizes to centromeres from mitosis to G1 Single optical section of S2 cells are shown. Boxes indicate the 3x enlarged inset. Scale bars represent 3µm. **a** Time-lapse live imaging pictures of cells transiently expressing Spt6-GFP and dCENP-A-mCherry. Numbers indicate minutes before/after onset of anaphase. **b** Single optical section of fixed early G1 cell expressing Spt6-GFP. dCENP-A immunodetection served as a marker of centromeres; tubulin staining allowed midbody detection.

**Supplemental Fig. 2.**
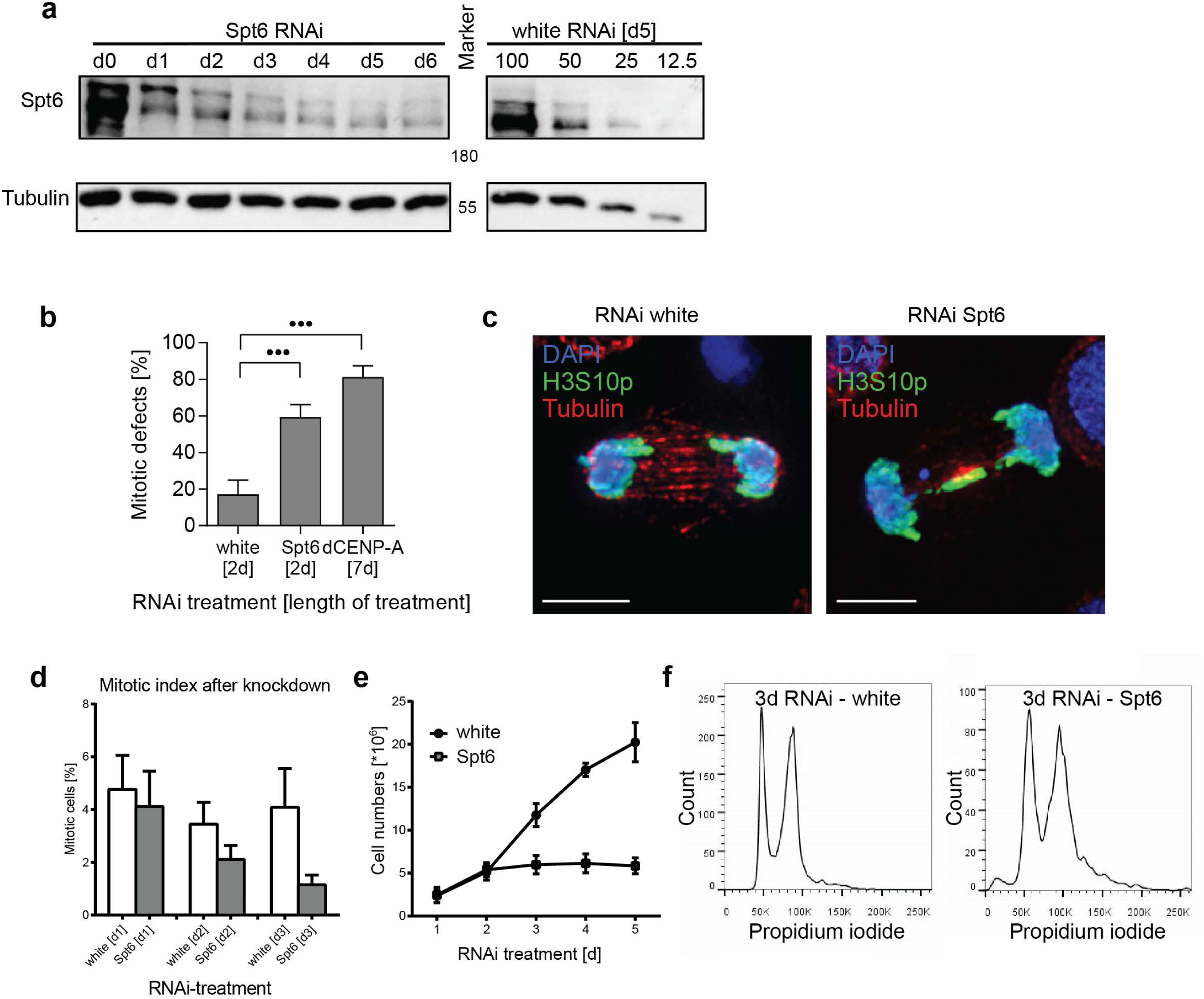
RNAi-mediated depletion of Spt6 results mitotic defects and in an unspecific cell-cycle block. **a** Western Blot analysis of Spt6 protein levels in whole cell lysates after RNAi treatment. White RNAi served as a control. Spt6 detection by Western Blot resulted in two bands (five different antibodies tested), with the upper band likely representing a modified version of Spt6. **b** Quantification of mitotic defects after white (control), Spt6 or dCENP-A RNAi. N=4 replicates; n=25 anaphase cells. Data are represented as mean+SD. • P≤0.05; •• P≤0.01; ••• P≤0.001; n.s.=not significant. **c** Maximum intensity projection of representative mitotic cells for white and Spt6 RNAi that were quantified in **b**. **d** Mitotic index after control (white) or Spt6 RNAi. Data are represented as mean+SD. N=3 replicates, n=500-700 cells. **e** 5d growth curve of S2 cells after RNAi-mediated depletion of control (white) or Spt6. N=3 replicates. Data are represented as mean+/-SD. **f** Single cell FACS profile of propidium iodide staining after 3d of white or Spt6 RNAi.

**Supplemental Fig. 3.**
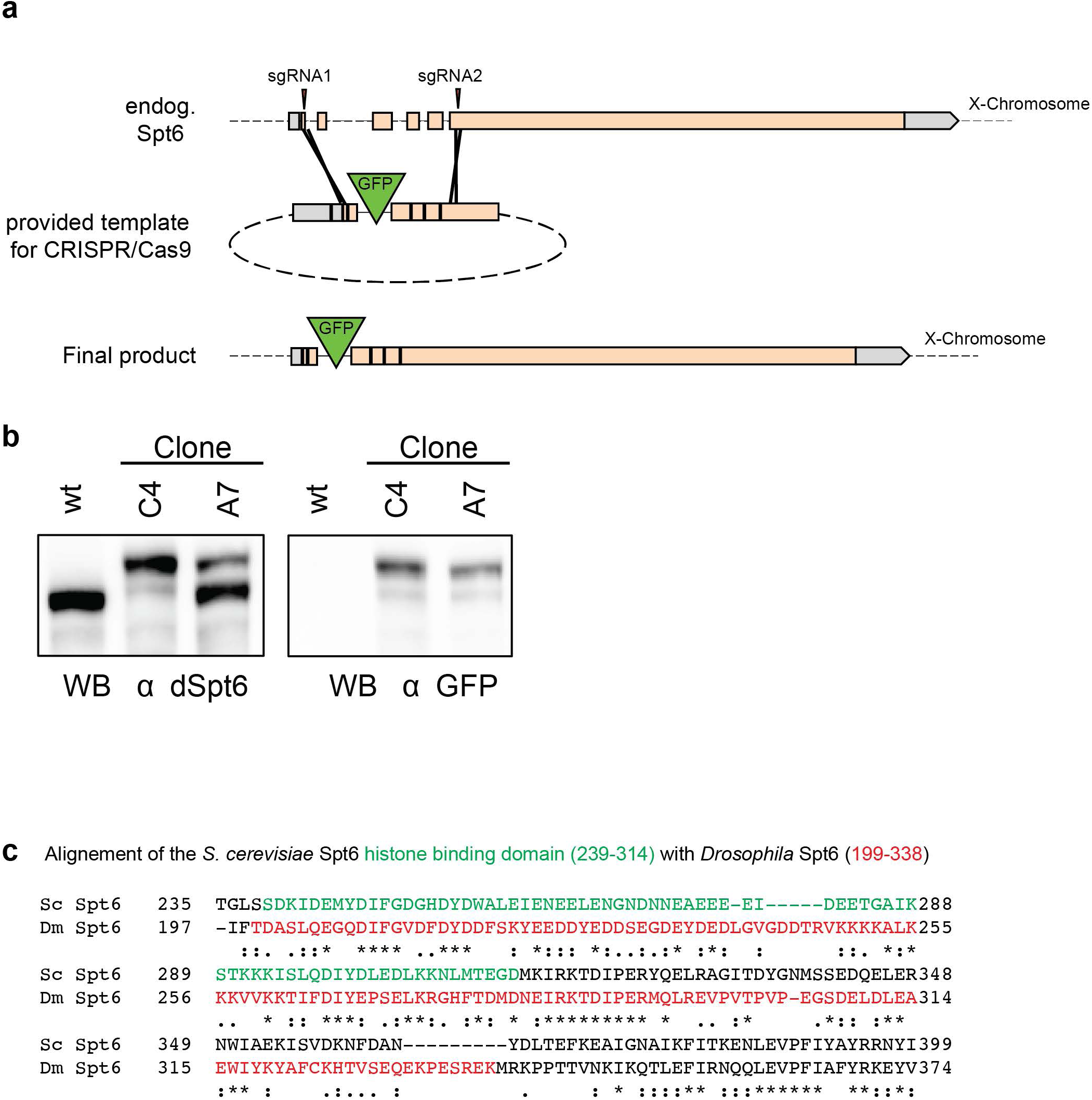
GFP-tagging of endogenous *Drosophila* Spt6 and alignment with the *S.c.* Spt6 histone binding domain. **a** Schematic representation of the GFP-Spt6 version created by CRISPR/Cas9. **b** Western blot analysis of Spt6 levels in whole cell extracts of wildtype (wt) cells and two clonal cell lines. Both alleles are tagged with GFP in clone C4, only one of the two genetic loci is tagged in clone A7. **c** Alignment of the histone binding domain of *S. cerevisiae* Spt6 (residues 239-314 shown in green) (McDonald et al. 2010) with the corresponding region of *Drosophila* Spt6. The bacterially expressed fragment of *Drosophila* Spt6 (199-338) is shown in green. Alignments were performed on Uniprot using the Clustal Omega program (Anon 2017). * (asterisk) = fully conserved residues, : (colon) = strongly similar residues,. (period) = weakly similar residues.

**Supplemental Fig. 4.**
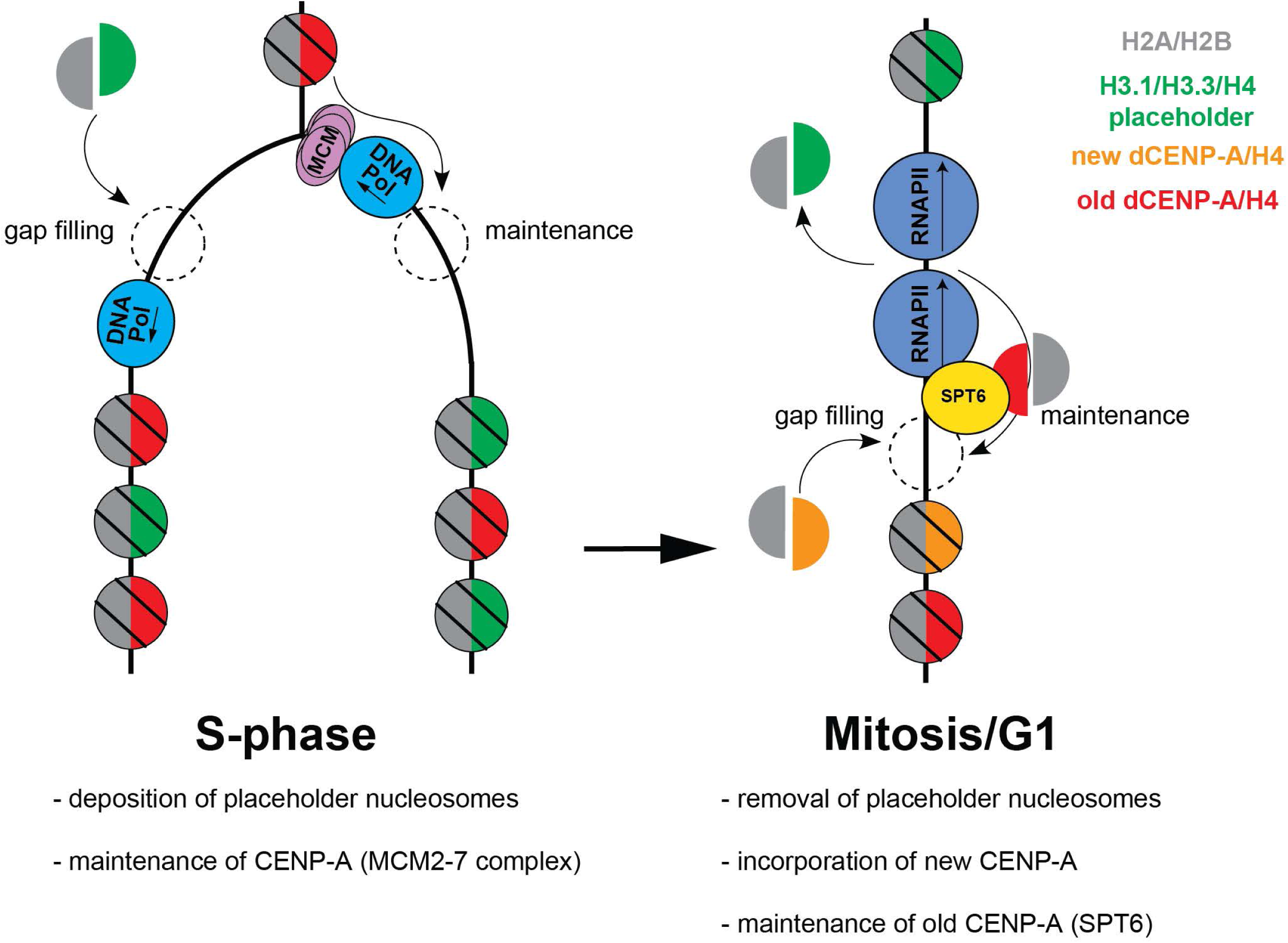
Schematic representation of nucleosome dynamics during the remodeling of centromeric chromatin in S-phase and Mitosis/G1.

